# Differentiating spermatogonia trigger somatic support cells to form septate junctions required for germ cell survival

**DOI:** 10.1101/2024.04.02.587826

**Authors:** Cameron W. Berry, Sarah R. Stern, Hannah Vicars, Lorenzo Gallicchio, Margaret T. Fuller

**Affiliations:** Department of Developmental Biology, Stanford University School of Medicine, USA; Department of Genetics, Stanford University School of Medicine, USA

**Keywords:** Septate Junctions, Somatic support cells, Germ cell survival, Spermatogenesis

## Abstract

Male germ cells from insects to mammals differentiate in a privileged microenvironment created by stage-specific junctional seals between somatic support cells that isolate meiotic and postmeiotic germ cells from bodily fluids. Here we show that action of the *Drosophila* germ cell differentiation factor Bag-of-marbles (Bam) is required for the somatic cyst cells associated with each cluster of transit amplifying spermatogonia to form the junctions that seal the cyst, isolating differentiating germ cells. Knockdown of septate junction (SJ) components or the transmembrane protein Side-V in somatic cyst cells resulted in elimination of most transit amplifying spermatogonia at the 8-cell stage. Germ cell death was spared in males mutant for *bam,* indicating that intact barriers surrounding transit amplifying progenitors are required to ensure germline survival only once differentiation has initiated. Together these results suggest that signals from spermatogonia initiating differentiation trigger their partner somatic cyst cells to form the tight junctional barrier. The close timing of onset of differentiation and the requirement for cyst cell sealing may account for the normal elimination of 20-30% of early germ cells prior to meiosis.

## Introduction

During animal development, cells arising from different embryonic origins come together to form functional organs, often delimited by one or more layers of cells arranged in an epithelium. In many cases the epithelial cells form occluding junctions with their neighbors, creating a barrier that seals off the internal cells of the organ from the rest of the body (Marchiando et al., 2010), or the tissue from compartments topologically connected to the outside, such as airways of the lung, intestinal lumen, or fluid contents of the bladder. As tissues form, critical cross check mechanisms eliminate malformed assemblies before they adversely affect tissue function resulting in the death of progenitor cells (Fuchs & Steller, 2011; Ghose & Shaham, 2020), observed in Rat oligodendrocytes (Barres & Raff, 1999), mammalian germ cells (Jeyaraj et al., 2003; Sinha Hikim & Swerdloff, 1999), and *Drosophila* germ cells (Yacobi-Sharon et al., 2013). This widespread cellular pruning may be especially important in systems where tissues must be replenished throughout life or repaired after injury by differentiation from adult stem cell founders.

We are investigating the formation and differentiation of *Drosophila* male germ line cysts as a model mini-organ containing a clone of sister germ cells enclosed by a simple squamous epithelium composed of two somatically derived cyst cells (Figure 1A). In *Drosophila*, as in mammals, male germ cells arise and differentiate in intimate contact with somatic support cells, which seal meiotic and postmeiotic germ cell stages off from the body. *Drosophila* spermatogonia and their enclosing somatic cyst cells arise from respective adult stem cells attached to a common niche at the apical tip of the testis (Fuller & Spradling, 2007). The male germ line stem cell (GSC) divides with an oriented spindle to produce a new stem cell and a daughter gonialblast (Gb), which is displaced away from the niche. The gonialblast initiates a series of four spermatogonial mitotic divisions with incomplete cytokinesis, producing a cyst containing 16 interconnected mitotic sister germ cells, which together enter premeiotic S phase then meiotic prophase. The somatic cyst stem cells (CySCs) also divide to both self renew and produce daughter cyst cells (CC) displaced away from the hub. Two somatic cyst cells flank and send out processes to enclose each gonialblast, pinching it off from its parent male germ line stem cell (Lenhart & DiNardo, 2015) and forming the male germ line cyst. The two cyst cells never divide again, but enclose and co-differentiate with the germ cells, expanding to cover all the progeny of the founding gonialblast throughout the entire process of spermatogenesis.

**Figure 1.**
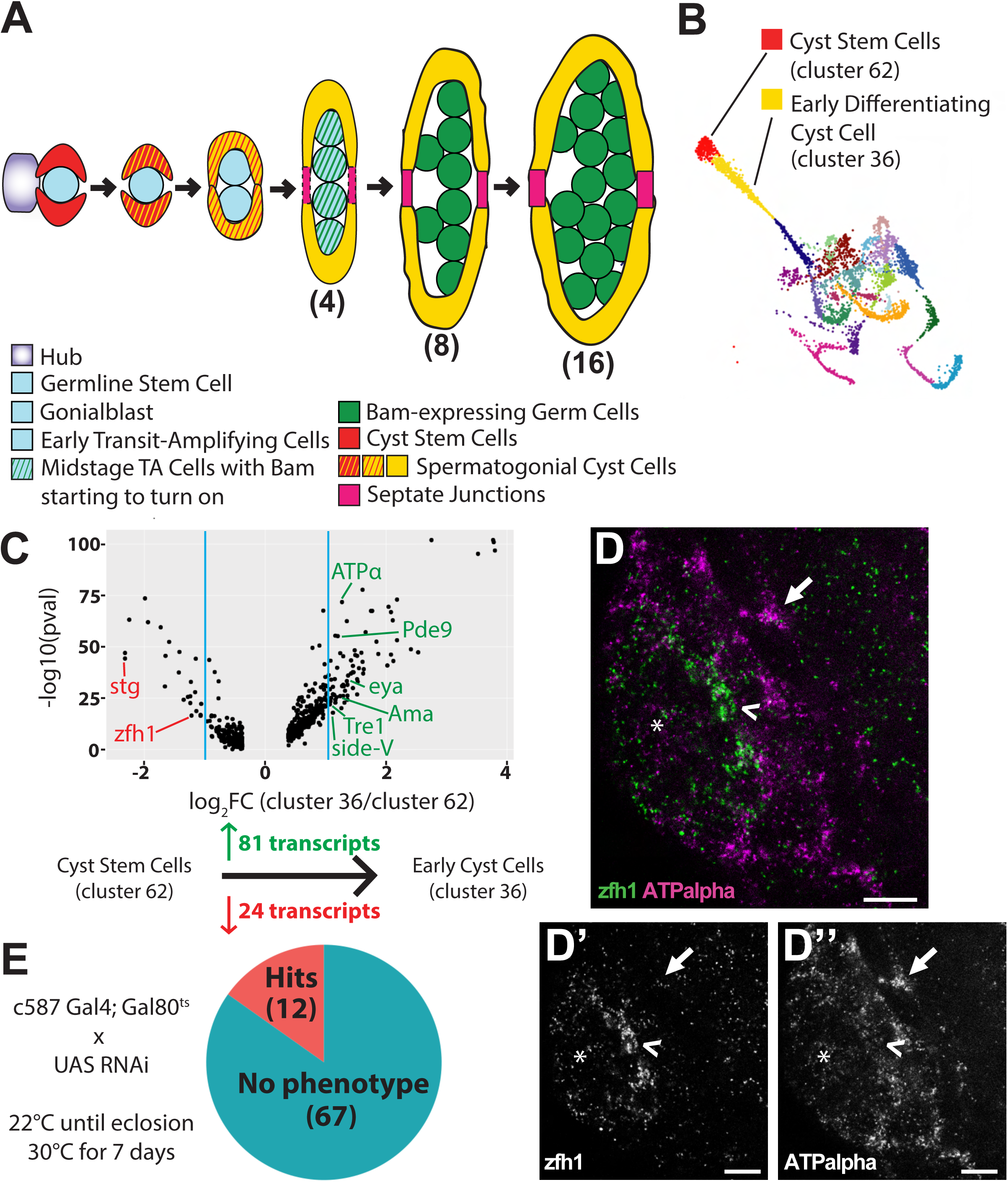
Genes upregulated at the earliest stages of cyst cell differentiation are required for proper spermatogenesis. (A) Diagram of early stages in *Drosophila* spermatogenesis depicting cyst cell and germline lineages as a germline stem cell differentiates into late transit amplifying cells. (B) UMAP plot of the cyst cell lineage generated from testis snRNAseq data (Li et al., 2022), colored by clusters generated by Leiden 6.0 clustering. (C) Volcano plot of genes differentially expressed between cluster 62 and cluster 36 with log_2_FC values greater than 1, generated with ASAP. (Below) Number of differentially expressed genes identified either as upregulated (green) or downregulated (red) between cluster 62 and cluster 36. (D) RNA Fluorescence *in situ* images of the apical region of testes stained with RNA FISH probes against (green) zfh1 and (magenta) Atpα in wild-type testes (n=5). (E) Fly crosses and temperature shift regimen employed to induce cyst cell-specific RNA knockdown of candidate genes. Pie chart indicating number of genes for which knockdown in cyst cells by RNAi produced a phenotype as assessed by phase-contrast microscopy of dissected testis tissue. Arrowhead: early differentiating cyst cell expressing high levels of zfh1; Arrow: differentiated cyst cell lacking zfh1 expression. Scale bars: 10 μm.

When the proliferating spermatogonia reach the 4-8 germ cell stage, septate junctions (SJs; insect occluding junctions) form between the two cyst cells to seal the cyst (Fairchild et al., 2015). Formation of this permeability barrier is important, as knock down of septate junction components in somatic cyst cells dramatically reduced production of spermatocytes and spermatids (Fairchild et al., 2015). Similarly, in mammalian testes, as spermatogonia begin to differentiate into spermatocytes, tight junctions between somatic Sertoli cells form a barrier that separates spermatocytes and spermatids from body fluids and creates a special compartment in which meiosis and spermiogenesis take place (Mruk & Cheng, 2015). Disruption of the Sertoli cell blood-testis barrier by loss of the tight junction protein Claudin-11 in mice resulted in loss of spermatocytes and male sterility (Gow et al., 1999; Mazaud-Guitot et al., 2010).

One outstanding question is what sets the timing of formation of the septate/tight junctions between cyst cells, so that cysts containing early spermatogonia are permeable, but later stage differentiating germ cells are properly sealed away from bodily fluids and circulating peptide signaling molecules. A key regulator of differentiation of spermatogonia toward spermatocyte state in *Drosophila* is expression in germ cells of the Bag of marbles (Bam) protein. In *bam* mutant males, spermatogonia fail to differentiate into spermatocytes but instead continue to proliferate for several extra rounds of mitosis, producing cysts containing 64, 128, or more spermatogonia enclosed in the two cyst cells. Normally, Bam protein begins to be detected by immunofluorescence microscopy around the 4-cell stage, with the signal increasing in intensity in 8-cell stage cysts. Gene dosage experiments indicated that onset of germ cell differentiation is trigged when Bam protein accumulates to a critical threshold (Insco et al., 2009).

Here we show that in absence of *bam* function, the cysts fail to seal, suggesting that onset of differentiation of germ cells may set the timing of septate junction formation by the overlying cyst cells. Furthermore, loss of function of septate junction components or the transmembrane cell surface protein Side-V in somatic cyst cells caused germ cells to die between the late 4-cell and early 8-cell stage, soon after onset of Bam protein expression. Notably, germ cells that lacked *bam* function survived and continued to proliferate as spermatogonia in testes where *side-V* or the SJ component *Neurexin-IV* (*Nrx-IV*) was knocked down in cyst cells.

Together our findings suggest a model that onset of germ cell differentiation (triggered by Bam) signals overlying cyst cells to form septate junctions that seal the cyst. However, if the septate junctions do not form promptly the differentiating germ cells die. The close timing of these events may account for the observation that 20-30% of late spermatogonia normally undergo germ cell death through a mechanism involving the Htra2 related serine protease Omi, involved in caspase-independent cell death (Yacobi-Sharon et al. 2013). Consistent with this, the germ cell death caused by knock down of *Nrx-IV* function in cyst cells was substantially alleviated by lowering the dosage of *omi*.

## Results

### Differential expression analysis of testis snRNAseq data reveals transcripts upregulated as cyst stem cells initiate differentiation

Differential expression analysis using ASAP (Gardeux et al., 2017) of clusters of cells from *Drosophila* testis snRNA seq data (Li et al., 2022) identified transcripts upregulated as cyst stem cells differentiate into early cyst cells. Cluster 62 was previously identified as containing cyst stem cells and cluster 36 as containing the earliest differentiating cyst cells (Figure 1B; Raz et al., 2023). ASAP identified 24 transcripts downregulated by at least 2-fold in cluster 36 compared to cluster 62 (Figure 1C). These included the mitotic marker *string* (*stg*) and the Jak/STAT-target *Zinc finger homeodomain protein 1 (zfh1)* required for cyst stem cell maintenance (Leatherman & Dinardo, 2008), consistent with the assignment of cluster 62 as cyst stem cells and cluster 36 as early differentiating cyst cells. Reciprocally, ASAP identified 81 transcripts upregulated more than 2-fold in cluster 36 compared to cluster 62, indicating transcription of these genes increases as cyst stem cells differentiate into early cyst cells (Figure 1C, Table S1). The 81 upregulated transcripts included *eyes absent* (*eya*), consistent with the increase in Eya protein observed in nuclei of cyst cells associated with transit amplifying spermatogonia and later stage cysts observed by immunofluorescence (Fabrizio et al., 2003). Fluorescent *in situ* hybridization (FISH) to wild-type testes confirmed increase in *Atpα* and decrease in *zfh1* mRNA levels in early differentiating cyst cells located a few cell diameters out from the apical hub (Figure 1D).

### Cell type-specific knock down of upregulated transcripts identifies genes required in cyst cells to limit germ cell proliferation

A cyst cell RNAi screen to knock down expression of transcripts upregulated as cyst stem cells differentiate into early cyst cells revealed both known and novel genes required in cyst cells for early steps in germ cell differentiation. To knock down gene function in cyst cells, flies carrying transgenes encoding *c587Gal4* and a temperature-sensitive Gal4 repressor, *tub-Gal80^ts^*, were crossed to fly lines carrying UAS-RNAi constructs targeting 79 of the 81 upregulated genes (Table S1). *c587Gal4* drives expression in the cyst cell lineage (Kai & Spradling, 2003), as well as in other somatic tissues throughout the fruit fly. The progeny were raised to adulthood at 22°C to allow normal testis development, then shifted to 30°C to initiate knock down (Figure 1E). Following 7 days of knock down at 30°C, testes were dissected and examined by phase contrast light microscopy to identify effects on germ cell differentiation. Testes from control flies carrying the *c587Gal4* expression driver and *tub-Gal80^ts^* but lacking the RNAi line and raised under the same temperature shift regimen resembled wild type, with small cells (dotted outline) at the apical tip including germline stem cells and transit amplifying spermatogonia, many larger cells in spermatocyte stages (bracket) further from the tip (Supplemental Figure 1A), and elongated spermatid bundles resulting from germ cell differentiation. Targeting 79 candidate genes with a total of 255 RNAi lines (Table S1) under these conditions identified 12 genes where knock down in the cyst cell lineage showed clear phenotypes when assessed by phase contrast light microscopy (Figure 1E, Supplemental Figure 1).

Knock down of *Ama*, *Tre1* or *Pde9* in cyst cells resulted in increased numbers of small cells at the tip of the testis compared to control, with no or few spermatocytes remaining after 7 days at 30°C (Supplemental Figure 1A-D). Immunofluorescence staining with anti-Vasa to mark germ cells confirmed that testes in which Pde9 was knocked down in the cyst cell lineage by RNAi had a large cluster of small germ cells at the apical tip and lacked cells expressing the spermatocyte marker LolaF (Supplemental Figure 1F). In control testes, staining for the mitotic marker phospho-histone H3 (pH3) only showed pH3 positive germ cells (marked by anti-Vasa) within a few cell diameters of the hub (Supplemental Figure 1G). In contrast, testes lacking Pde9 function in cyst cells showed pH3 positive germ cells in mitosis much farther from the hub. These were commonly singlets, doublets, or only very small clusters, indicating that the germ cells were proliferating as gonialblasts or early spermatogonia (Supplemental Figure 1H), rather than as large cysts of germ cells undergoing synchronized divisions as in *bam* mutant males.

### Function of septate junction (SJ) components and *side-V* is required in somatic cyst cells for survival of late transit amplifying spermatogonia

In contrast, knock down of the septate junction (SJ) component *Atpα* or the motor neuron axon guidance receptor *side-V* (Sink et al., 2001) in the cyst cell lineage resulted in testes with only a few small cells remaining at the testis tip (Figure 2B, C), compared to control testes, which contained many germ cells at different stages of differentiation (Figure 2A). Under the knock down conditions assessed, loss of function of *side-V* or *Atpα* in the cyst cell lineage resulted in testes with elongated spermatid bundles (from germ cells that had started differentiation before the temperature shift), but lacking spermatocytes and early spermatid stages. Notably, a limited number of small cells remained clustered at the apical tip.

**Figure 2.**
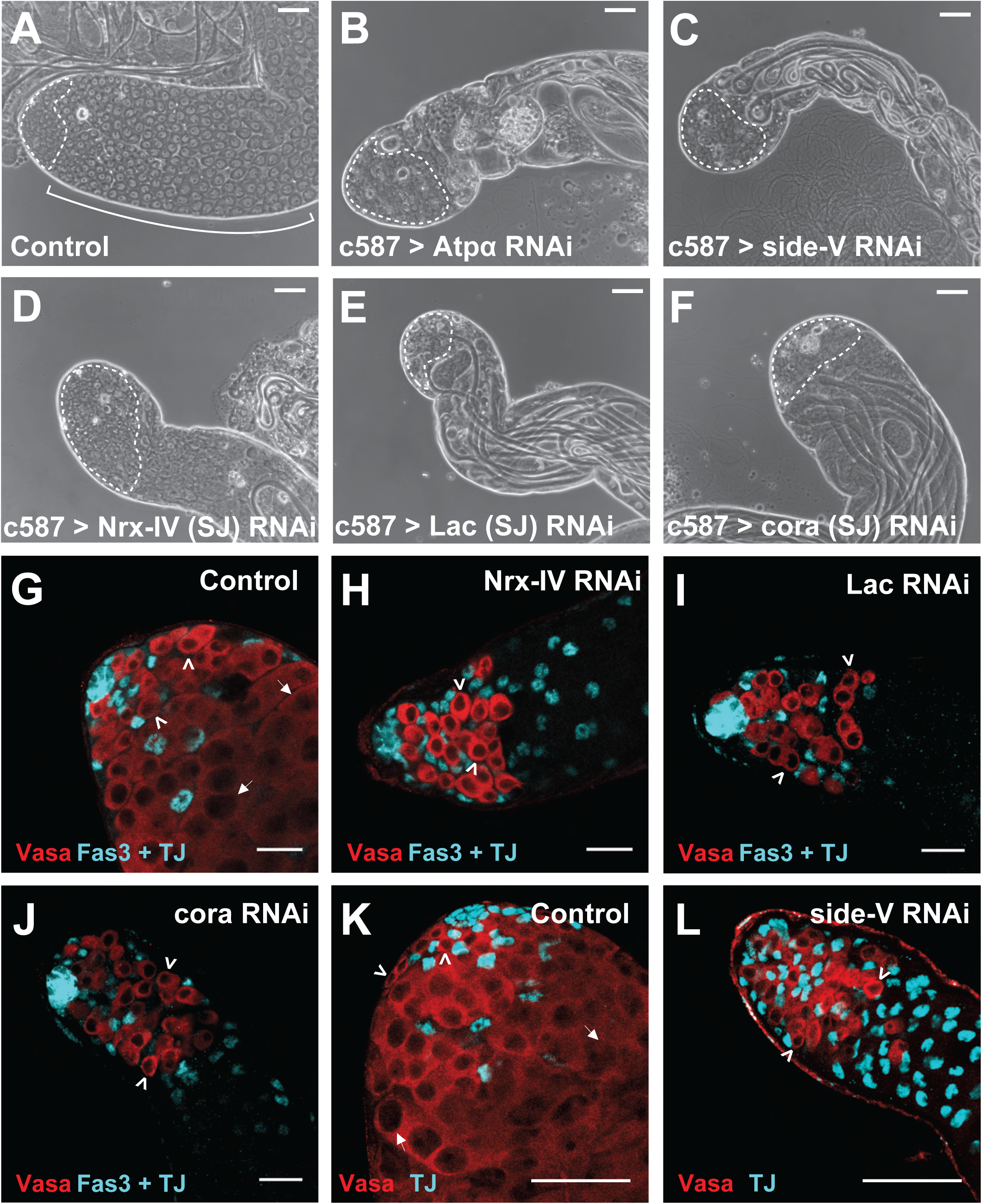
Cyst cell RNAi of septate junction components or *side-V* leads to loss of differentiating germ cells. (A-F) Phase-contrast images of the apical region of testes from male *c587Gal4, Gal80^ts^* flies shifted to 30°C for 7 days that had (A) no RNAi transgene (n=10) or an RNAi transgene targeting (B) *Atpα* (n=7), (C) *side-V* (n=11), (D) *Nrx-IV* (n=9), (E) *Lac* (n=5) or (F) *cora* (n=7). (G-J) Immunofluorescence images of the apical region of testes stained with (teal) anti-Fas3 and anti-TJ and (red) anti-Vasa antibodies from male *c587Gal4*, *Gal80^ts^* flies shifted to 30°C for 7 days that had (G) no RNAi transgene (n=15), or an RNAi transgene targeting (H) *Nrx-IV* (n=13), (I) *Lac* (n=13), or (J) *cora* (n=12). (K-L) Immunofluorescence images of the apical region of testes stained with (teal) anti-TJ and (red) anti-Vasa antibodies from male *c587Gal4, Gal80^ts^* flies shifted to 30°C for 7 days that had (K) no RNAi transgene (n=11), or an RNAi transgene targeting (L) *side-V* (n=7). Dotted lines, arrowheads: spermatogonia; Curved bracket, arrows: spermatocytes;: spermatid bundles. Scale bars: 50 μm.

Further investigation showed that knock down in cyst cells of several other septate junction components, including *Neurexin-IV* (*Nrx-IV*), *Lachesin* (*Lac*), and *coracle* (*cor*), resulted in phenotypes similar to knock down of *Atpα*, assessed by phase contrast microscopy (Figure 2D-F), suggesting that canonical septate junctions in cyst cells play an important role in supporting survival of germ cells in the late spermatogonial transit amplifying divisions. Immunofluorescence staining with anti-Vasa to mark germ cells and anti-TJ to mark the nuclei of cyst stem cells and early cyst cells showed that control testes from flies carrying *c587Gal4* and *tub-Gal80^ts^* but lacking the RNAi construct subjected to the knock down temperature regime had abundant early germ cells and spermatocytes (Figure 2G). In contrast, by 7 days after the shift to 30°C to initiate knock down, loss of function of the core transmembrane SJ components *Neurexin-IV* (*Nrx-IV*) or *Lachesin* (*Lac*) or of the cytoplasmic SJ component *coracle* (*cor*) in the cyst cell lineage resulted in the loss of early germ cells undergoing spermatogonial divisions. In all three cases, cyst cell knockdown of the septate junction components resulted in testes that retained a small group of germ cells at the tip of the testis (Figure 2H-J) but lacked spermatocytes. Knock down of *side-V* in cyst cells also led to loss of spermatocytes, but not TJ-positive cyst cells (Figure 2L) compared to control testes (Figure 2K). Strikingly, most testes did not show overproliferating germline cysts following knock down of either *side-V* or the septate junction component *Nrx-IV*.

Analysis of time points soon after induction of RNAi revealed that late spermatogonia were lost to cell death when function of the SJ component *Nrx-IV* was knocked down in somatic cyst cells. An increase in the number of TUNEL-positive cells compared to controls appeared as early as 2 days post induction of RNAi against *Nrx-IV* in cyst cells (Figure 3A, B). The TUNEL-positive cells were located in a zone of germ cell death in the region where late transit amplifying spermatogonial germ cells normally reside. Testes observed at later time points after *Nrx-IV* knockdown also had increased TUNEL staining at the testis tip compared to controls raised with the same temperature regimen. At day 3 post onset of *Nrx-IV* knockdown, large, maturing spermatocytes were adjacent to spermatogonia (Figure 3C), suggesting that germ cells beyond the zone of cell death, presumably generated prior to the knockdown, continued to differentiate. Prolonged knockdown resulted in morphological changes, including a thinner testis shape (Figure 3E), likely due to decline in the number of differentiating spermatocytes. By day 5 of *Nrx-IV* knockdown (Figure 3G) testes resembled day 7 of *Nrx-IV* knockdown (Figure 2H), with a small group of germ cells remaining at the testis tip. Similarly, knock down of *side-V* in cyst cells resulted in large, maturing spermatocytes adjacent to spermatogonia by day 3 (Figure 3K) and a decrease in testis width compared to controls after several days of knock down (Figures 2C; 3M-P).

**Figure 3.**
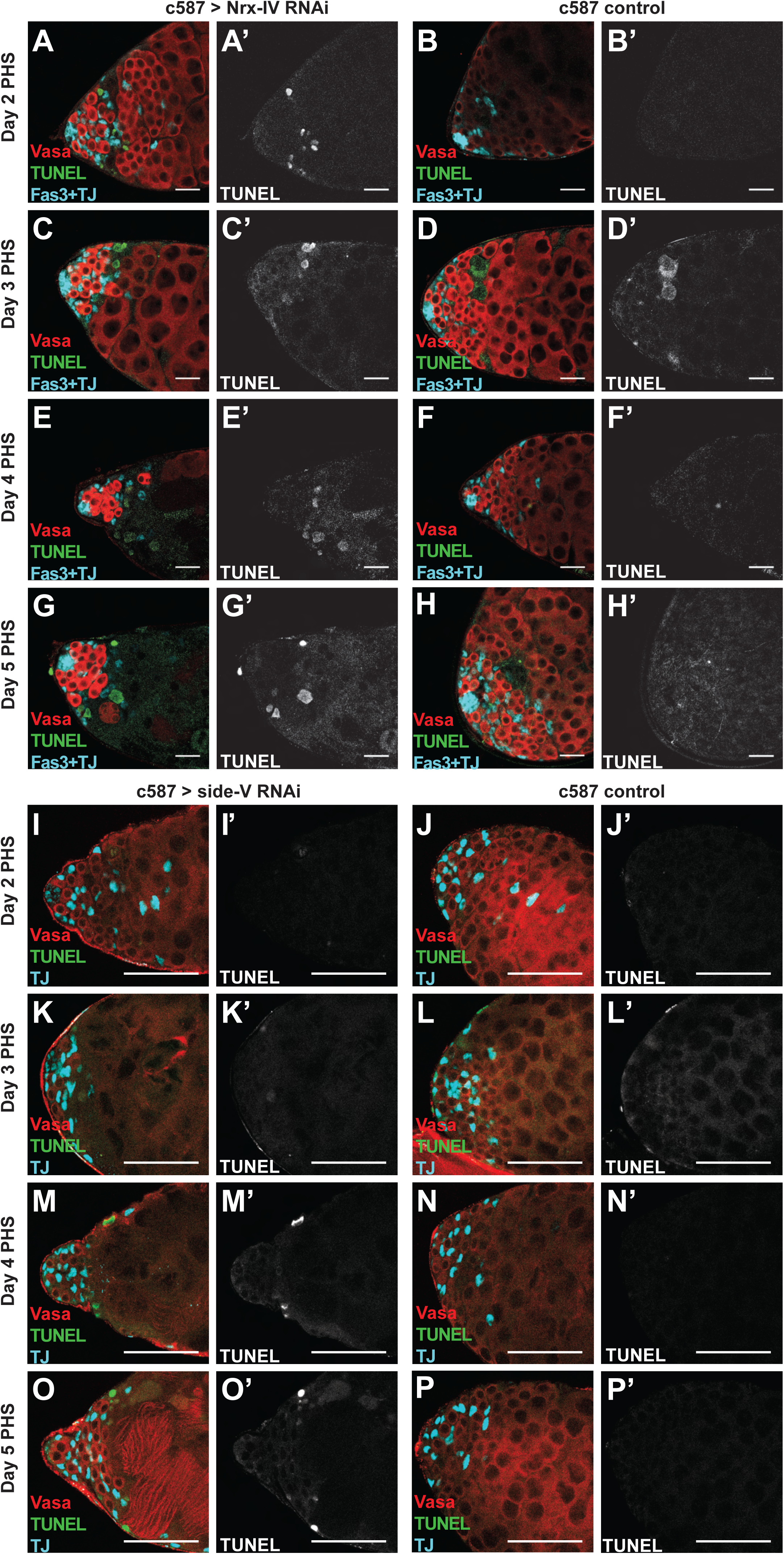
Loss of *Nrx-IV* or *side-V* in cyst cells results in acute germ cell death at mid transit amplifying stages. (A-H) Immunofluorescence images of the apical region of testes stained with (teal) anti-Fas3 and anti-TJ, and (red) anti-Vasa antibodies along with (green/white) TUNEL from male *c587Gal4, Gal80^ts^*flies shifted to 30°C for the specified number of days. Control flies with no RNAi construct were shifted to 30°C for (B) 2 (n=6), (D) 3 (n=5), (F) 4 (n=5) or (H) 5 (n=3) days prior to dissection. *Nrx-IV* knockdown flies, which had the RNAi transgene targeting *Nrx-IV*, were shifted to 30°C for (A) 2 (n=4), (C) 3 (n=5), (E) 4 (n=5) or (G) 5 days (n=5) prior to dissection. (I-P) Immunofluorescence images of the apical region of testes stained with (teal) anti-TJ and (red) anti-Vasa antibodies along with (green/white) TUNEL from male *c587Gal4, Gal80^ts^* flies shifted to 30°C for the specified number of days. Control flies with no RNAi construct were shifted to 30°C for (J) 2 (n=11), (L) 3 (n=3), (N) 4 (n=6) or (P) 5 (n=8) days prior to dissection. *side-V* knockdown flies, which had the RNAi transgene targeting *side-V*, were shifted to 30°C for (I) 2 (n=3), (K) 3 (n=10), (M) 4 (n=7) or (O) 5 days (n=12) prior to dissection. Scale bars: 50 μm.

### *Nrx-IV* is required in the cyst cell lineage for germ cells to survive beyond the 4-cell transit amplifying stage

Staining of ring canals and fusome to allow precise quantification and analysis of the stage of transit amplifying germ line cysts revealed that early spermatogonial cysts were present in normal numbers after knockdown of *Nrx-IV* in somatic cyst cells, but that the majority of spermatogonia did not survive beyond the 4-cell stage. During the transit amplifying spermatogonial divisions, germ cells undergo incomplete cytokinesis, resulting in cysts of interconnected germ cells that are mitotic sisters, connected by ring canals. The fusome, a cytoplasmic, membranous structure, runs through these ring canals within a germline cyst, grouping the germ cells together in a cluster (Hime et al., 1996). The germ cells in 2-cell stage cysts have one ring canal strung on a short fusome connecting the two germ cells (Figure 4B). 4-cell cysts have 3 ring canals strung along a curved fusome (Figure 4C, 4H). Cysts at the 8- and 16-cell stages have 7 and 15 ring canals respectively, linked by a branched fusome.

**Figure 4.**
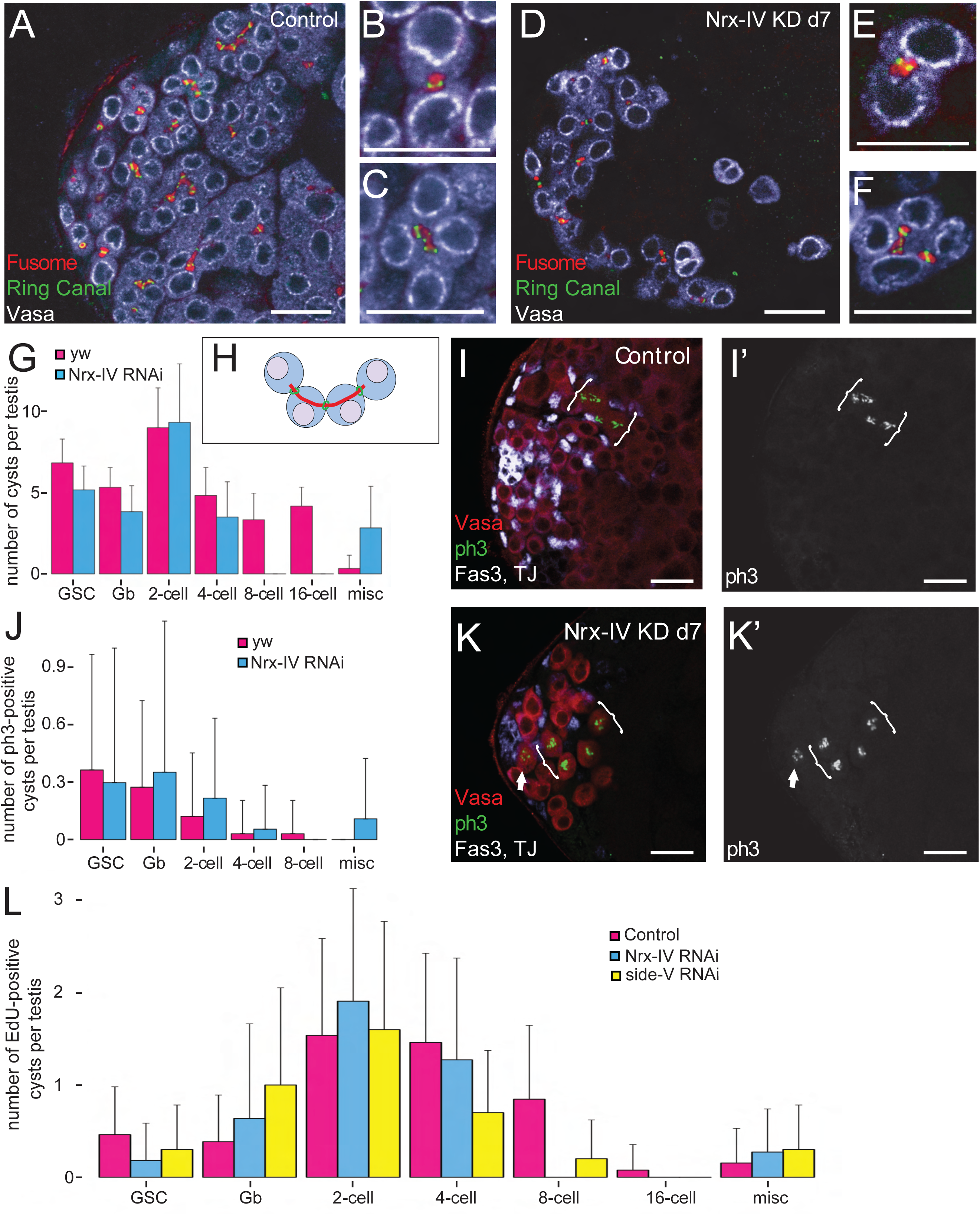
*Nrx-IV* is required in cyst cells for germ cells to progress beyond the 4-cell stage. (A, B) Immunofluorescence images of the apical region of testes stained with (red) anti-Hts, (green) the ring canal marker anti-cindr, and (white) anti-Vasa antibodies from male *c587Gal4, Gal80^ts^* flies shifted to 30°C for 7 days that had (A) no RNAi transgene or (D) an RNAi transgene targeting *Nrx-IV*. Higher magnification pictures show (B and E) 2-cell and (C and F) 4-cell spermatogonial cysts. (G) Quantification of the number of germline stem cell (GSC), gonialblast, 2-cell, 4-cell, 8-cell, 16-cell or miscellaneous cysts per testis (n=7 for yw; n=6 for *Nrx-IV* RNAi) and (J) quantification of the number of pH3-positive cysts per testis (n=33 for yw; n=36 for *Nrx-IV* RNAi) from *c587Gal4, Gal80^ts^* males shifted to 30°C for 7 days with (pink) no RNAi transgene or (blue) an RNAi transgene targeting *Nrx-IV*. (H) Diagram of a 4-cell transit amplifying cyst with fusome in green and ring canals in red. (I, K) Immuno-fluorescence images of the apical region of testes stained with (white) anti-Fas3 and anti-TJ, (red) anti-Vasa and (green/white) anti-pH3 antibodies from *c587Gal4, Gal80^ts^* male flies shifted to 30°C for 7 days that had (I) no RNAi transgene or (K) an RNAi transgene targeting *Nrx-IV*. (L) Quantification of the number of EdU-positive cysts per testis (n=13 for Control; n=11 for *Nrx-IV* RNAi; n=10 for *side-V* RNAi). Arrow: dividing germline stem cell; Brackets: cysts of mitotic 4-cell transit amplifying cells. Scale bars: 50 μm.

Immunofluorescence staining of ring-canal and fusome components in testes in which function of *Nrx-IV* had been knocked down in the somatic cyst cell lineage for 7 days revealed a lack of cysts with 8 or 16 germ cells. 4-cell cysts were as common in the knock down as in control testes subjected to the same temperature shift regimen (Figure 4G). However, many 4-cell cysts in *Nrx-IV* knock down testes had aberrant fusomes that did not correctly span all three ring canals (Figure 4D). In the 6 *Nrx-IV* knock down testes analyzed by ring canal-fusome staining, no 8-cell cysts were identified (Figure 4G), whereas twenty 8-cell cysts were identified in the 6 control testes stained in parallel. Additionally, testes in which *Nrx-IV* had been knocked down in the cyst cell lineage for 7 days contained germ cells that did not resemble germ cells observed in control testes, including a small percentage of escaper cysts that contained more than 16 cells (i.e. a 22-cell cyst) and isolated pairs of germ cells that had more than one ring canal, suggesting the two cells may have broken off from a 4-cell cyst. Staining for the mitotic chromosome marker pH3 to score cysts of germ cells undergoing synchronous mitosis indicated that spermatogonia up to the 4-cell stage were able to enter the mitotic division program prior to undergoing cell death under conditions where Neurexin-IV had been knocked down in somatic cyst cells. Quantification of germ cells undergoing mitotic division based on immunofluorescence staining with anti pH3 showed GSCs, Gbs, and germ cells in 2-cell or 4-cell cysts undergoing mitosis in testes where *Nrx-IV* had been knocked down in the cyst cell lineage, as in controls (Figure 4I-K). However, consistent with the lack of 8-cell cysts indicated by the analysis of ring canals and fusome above, no cysts with 8 germ cells in mitosis were detected in the 37 *Nrx-IV* knock down testes examined (Figure 4J). Similarly, labeling of germ cells undergoing DNA replication by brief incubation of testes in EdU revealed a striking decrease in 8-cell stage spermatogonial cysts undergoing S-phase when cyst cells lacked *Nrx-IV* compared to control testes (Supplemental Figure 2A,B; Quantified in Figure 4L). Knockdown of *side-V* in cyst cells also resulted in a normal number of 2-cell cysts in S-phase but a reduced number of 8-cell cysts with germ cells in S-phase compared to control testes labelled under the same conditions. The effect appeared more gradual than for knock down of *Nrx-IV*, with the number of cysts with 4 germ cells in S-phase also reduced (Figure 4L; Supplemental Figure 2C, D).

### Cyst cells persist in the absence of *Nrx-IV*, but activate the JNK pathway

Loss of *Nrx-IV* did not grossly affect the ability of cyst stem cells to self-renew or differentiate into early cyst cells. While *Nrx-IV* knockdown in the cyst cell lineage under control of *c587-Gal4* led to loss of germ cells, immunofluorescence staining revealed that cyst stem cells near the apical tip of testes in which function of *Nrx-IV* had been knocked down by RNAi under control of *c587Gal4* robustly expressed Zfh1, a readout of the response to Jak-Stat signaling characteristic of cyst stem cells and (at a reduced level in) cyst cell nuclei associated with proliferating spermatogonial cysts in wildtype (Supplemental Figure 3A, B). Immunofluorescence staining also confirmed that cyst stem cells and cyst cells lacking *Nrx-IV* near the apical tip of the testis expressed the nuclear marker *traffic jam (tj)*, with the level of Tj signal in cyst cell nuclei decreasing further from the apical tip, consistent with differentiation. Also consistent with cyst cell differentiation, *eyes absent (eya*) protein was detected in nuclei of cyst cells away from the testis tip, as in controls (Supplemental Figure 3A, B).

Although, based on the markers Eya and Tj, somatic cyst cells persisted after the death of late spermatogonia in *Nrx-IV* knockdown testes (Supplemental Figures 2B and 3B), the cyst cells remaining after germ cell death expressed high levels of *pucLacZ*, a readout of JNK pathway activation not normally expressed at high levels in cyst cells (Supplemental Figure 3C, D). Knock down of *side-V* similarly resulted in the persistence of cyst cells with increased puc-LacZ in the absence of their germ cell neighbors (Supplemental Figure 3E, F).

### Death of transit amplifying spermatogonia due to loss of *Nrx-IV* or *side-V* function in cyst cells requires the germ line differentiation factor Bam

The time at which spermatogonia began to show abnormalities (4-cell cysts) and then die (by the 8-cell cyst stage) in *Nrx-IV* knock down testis corresponded to when the germ cell differentiation factor Bag-of-marbles (Bam) becomes expressed. In wildtype testes, Bam protein begins to be detected by immunofluorescence staining in 4-cell cysts, with levels detected increasing in 8-cell stage spermatogonia (Figure 5A). Insco et al. (2009) showed that immunodetection of Bam protein remained high in early 16-cell cysts, then the protein abruptly disappeared after completion of premeiotic S phase. In testes in which *Nrx-IV* was knocked down in cyst cells, some germ line cysts at the edge of the death zone showed immunofluorescence signal for Bam protein, but the number of Bam positive cysts per testis was always low (Figure 5B).

**Figure 5.**
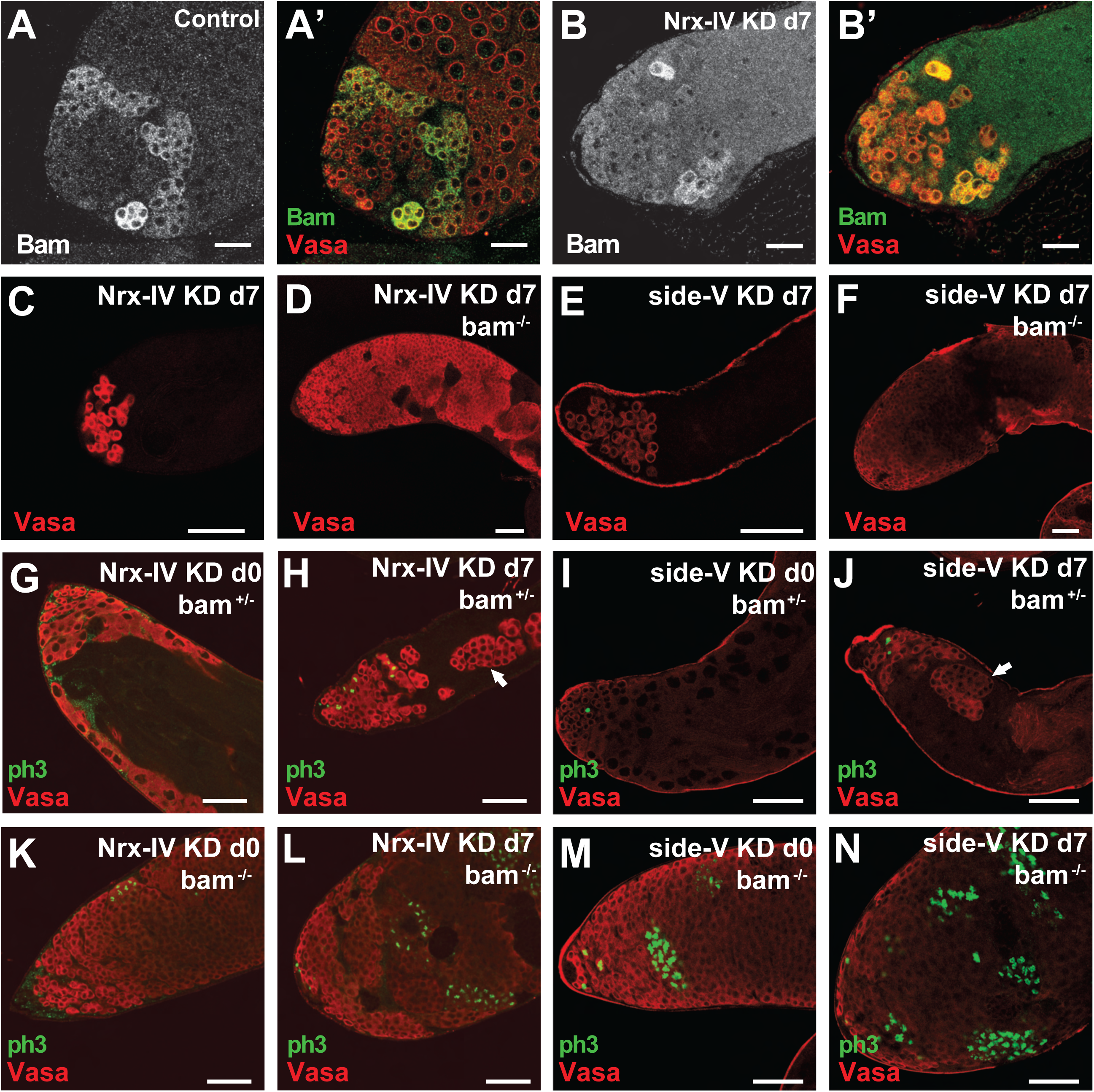
Bam is required for loss of *Nrx-IV* or *side-V* function in cyst cells to trigger transit amplifying germ cell death. (A, B) Immunofluorescence images of the apical region of testes stained with (red) anti-Vasa and (green) anti-Bam antibodies from male *c587Gal4, Gal80^ts^* flies shifted to 30°C for 7 days that had (A) no RNAi transgene (n=4) or (B) an RNAi transgene targeting *Nrx-IV* (n=5). (C, D) Immunofluorescence images of the apical region of testes stained (red) anti-Vasa from male *c587Gal4, Gal80^ts^* flies that had an RNAi transgene targeting *Nrx-IV* and were shifted to 30°C for 7 days that were (C) wildtype for *bam* (n=9) or (D) homozygous *bam*^−/−-^ (n=7). (E, F) Immunofluorescence images of the apical region of testes stained with (red) anti-Vasa antibody from male *c587Gal4, Gal80^ts^* flies that had an RNAi transgene targeting *side-V* and were shifted to 30°C for 7 days that were (E) wildtype for *bam* (n=5) or (F) homozygous *bam*^−/−-^ (n=12). (G-N) Immunofluorescence images of the apical region of testes stained with (red) anti-Vasa and (green) anti-pH3 antibodies from male *c587Gal4, Gal80^ts^* flies carrying an RNAi transgene targeting (G, H, K, L) *Nrx-IV* shifted to 30°C for (G, K) 0 as control or (H, L) 7 days that were (G, H) heterozygous *bam*^+/−^ (n=6, 8 respectively) or (K, L) homozygous *bam*^−/−-^ (n=8, 8 respectively), or carrying an RNAi transgene targeting (I, J, M, N) *side-V* shifted to 30°C for (I, M) 0 as control or (J, N) 7 days that were (I, J) heterozygous *bam*^+/−^ (n=5, 8 respectively) or (M, N) homozygous *bam*^−/−-^ (n=13, 10 respectively). Arrow: large cyst of spermatogonia. Scale bars: 50 μm.

The death of mid-stage transit amplifying spermatogonia following knock down of *Nrx-IV* or *side-V* in cyst cells appears to depend on the expression of Bam in the germline. Testes from males mutant for *bam* in which *Nrx-IV* or *side-V* was knocked down in cyst cells by RNAi had a large number of proliferating spermatogonia, in contrast to the small number of early germ cell cysts in males with *Nrx-IV* or *side-V* knock down alone (Figure 5C-F). Even a partial decrease in Bam levels improved germ cell survival (Figure 5G-J). A moderate increase in the number of germ cells was observed in testes when *Nrx-IV* or *side-V* was knocked down by RNAi in *bam^+/−^*heterozygotes compared to *Nrx-IV* or *side-V* knock down in flies with two wild type *bam* alleles (compare Vasa positive cells in Figures 5C and 5H, and Figures 5E and 5J), with proliferating escaper cysts observed beyond the group of germ cells at the tip of the testis (Figure 5H, J). Immunofluorescence staining for pH3 in *bam^−/−-^* homozygotes following *Nrx-IV* or *side-V* knock down revealed large testes filled with cysts of small germ cells which, based on pH3 staining, were undergoing mitosis in synchrony (Figure 5K-N).

### Cyst cells fail to form a permeability barrier in *bam^−/−-^* testes

Strikingly, incubation of testes in 3kDa or 10KDa dextran revealed that permeability barriers failed to form in *bam^−/−-^* testes. Consistent with the findings of Fairchild *et al*. (2015), 10 kDa fluorescent Dextran outlined individual GSCs, but was excluded from late transit amplifying spermatogonial cysts and all later stages of germ cell differentiation in testes wild type for *bam* or in most cysts in *bam^+/−^* heterozygotes. Similar results were obtained with 3 kDa dextran (Figure 6A, B, D, E). However, the cysts containing overproliferating spermatogonia in *bam^−/−-^* testes failed to exclude 3 kDa or 10 kDa Dextran (Figure 6C, F), indicating lack of permeability barrier formation, even though cyst cell membranes surrounded the clusters of overproliferating spermatogonia based on appearance of membrane-targeted mCD8-GFP expressed in cyst cells from a transgene under control of *c587-GAL4* (Figure 6C’, F’).

**Figure 6.**
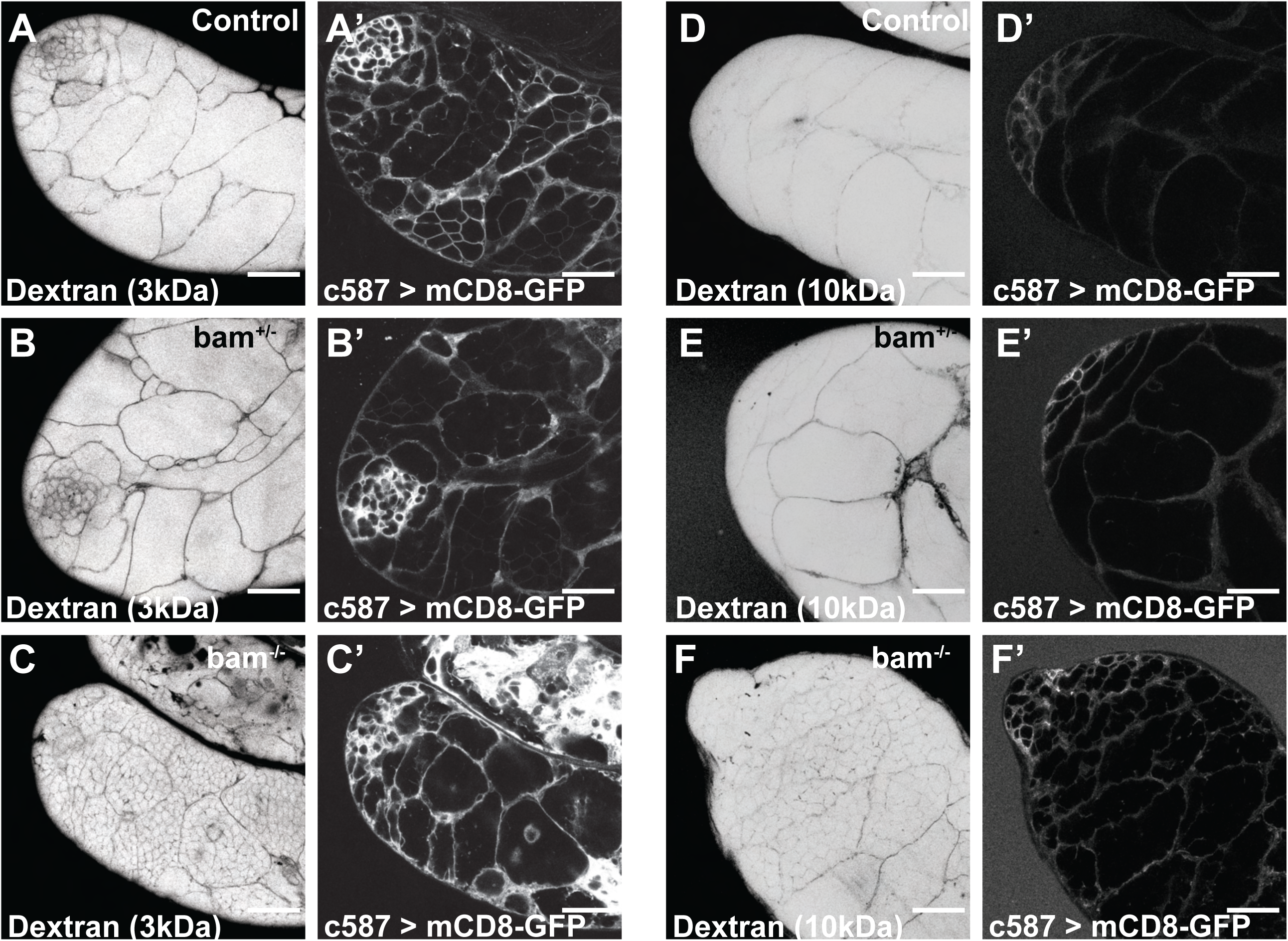
Cyst cells do not successfully form a permeability barrier around germ cells in testes lacking Bam. (A-C) Live permeability assay visualizing the apical region of testes incubated in (black, due to image inversion) fluorescently labeled 3 kDa Dextran and accompanying (A’-C’) live fluorescence images of (white) membrane-localized GFP in male *c587Gal4, UAS-mCD8-GFP* flies that were (A, A’) wildtype for *bam* (n=6) or (B, B’) heterozygous *bam^+/−^* (n=3) or (C, C’) homozygous *bam^−/−-^*(n=2). (D-F) Live permeability assay utilizing 10 kDa fluorescently labeled Dextran, under the same conditions as described for (A-C), in male *c587Gal4, UAS-mCD8-GFP* flies that were (D, D’) wildtype for *bam* (n=11) or (E, E’) heterozygous *bam*^+/−^ (n=5) or (F, F’) homozygous *bam*^−/−-^ (n=8). Scale bars: 50 μm.

### Germ cell death following knock down of *Nrx-IV* in cyst cells depends on *omi*

Death of spermatogonia in 8-cell cysts caused by loss of function of *Nrx-IV* in cyst cells likely occurred by the alternative germ cell death pathway that normally prunes spermatogonial cysts during homeostasis. Germ cell death following knock down of *Nrx-IV* in cyst cells was suppressed by lowered levels of *omi* (Figure 7), a component of an alternative cell death pathway known to contribute to death of *Drosophila* proliferating spermatogonia (Yacobi-Sharon et al., 2013). Strikingly, testes from *omi/+* males in which the function of *Nrx-IV* had been knocked down in cyst cells were filled with germ cells that remained small, indicating lack of differentiation (Figure 7B-E). The cyst cell nuclei associated with these rescued spermatogonia remained positive for the early cyst cell marker Tj (Figure 7B), indicating that they too did not progress to differentiation. Staining for the mitotic marker pH3 indicated that the rescued germ cells divided primarily in small clusters (Figure 7C) consistent with a mixed identity of gonialblasts and early transit amplifying cells, but did not show large clusters of synchronously dividing germ cells. Immunofluorescence staining for Bam revealed that most of the overproliferating germ cells were Bam negative (Figure 7D, E) with a small number of Bam positive germline clusters scattered down the testis. In contrast, in *omi*/+ testes in which *Nrx-IV* had not been knocked down, Bam positive cells were restricted to a ring near the apical tip of the testis (Figure 7D), as occurs in wildtype.

**Figure 7.**
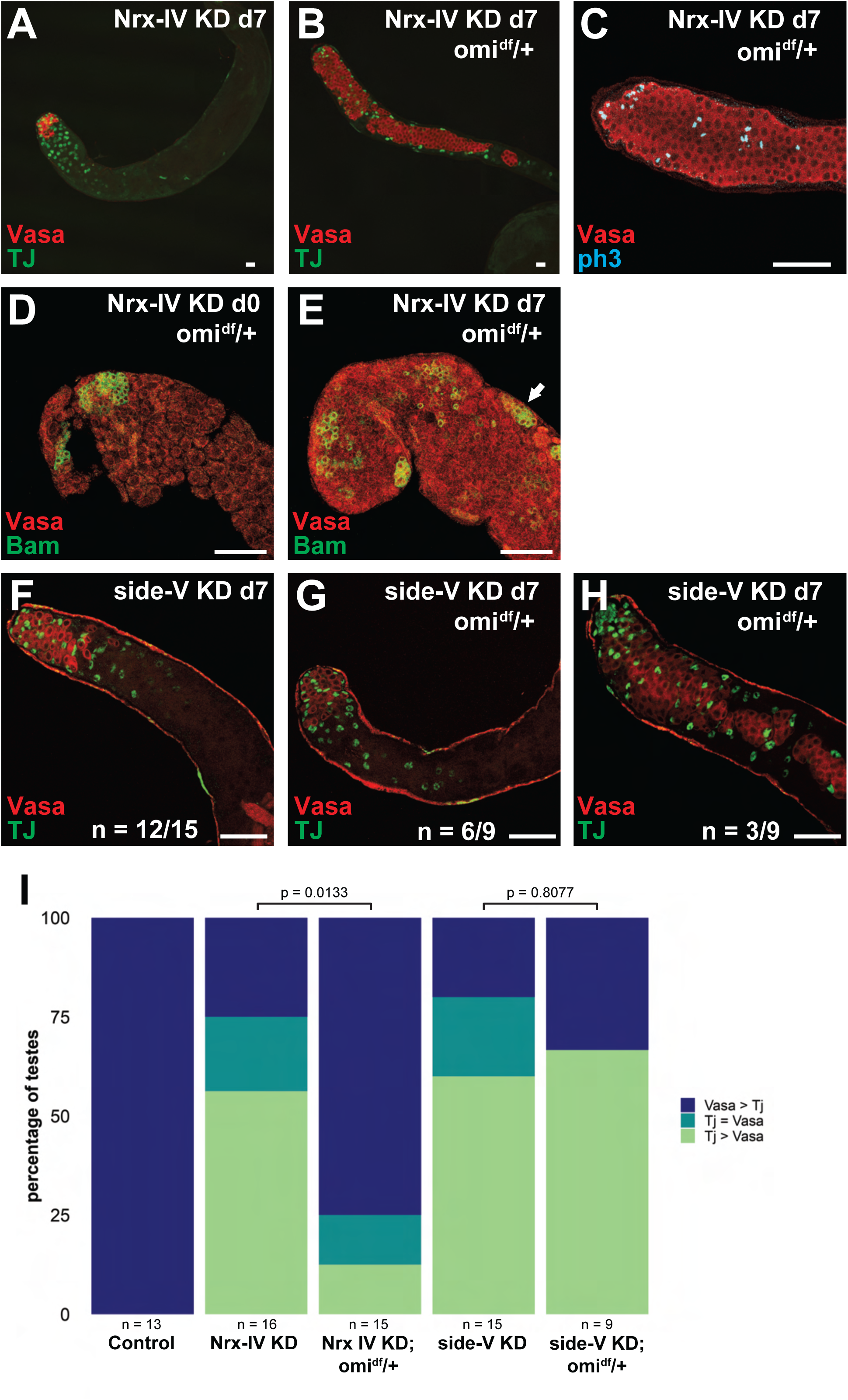
Germ cell death trigged by loss of *Nrx-IV* function in cyst cells depends on omi. (A, B) Immunofluorescence images of the apical region of testes stained with (red) anti-Vasa and (green) anti-TJ antibodies from male *c587Gal4, Gal80^ts^*flies shifted to 30°C for 7 days that had an RNAi transgene targeting *Nrx-IV* with (A) wild-type *omi* alleles (n=7) or (B) heterozygous *omi^df^*/+ (n=12). (C) Immunofluorescence images of the apical region of testes stained with (red) anti-Vasa and (teal) anti-pH3 antibodies from male *c587Gal4, Gal80^ts^* flies shifted to 30°C for 7 days that had an RNAi transgene targeting *Nrx-IV* with heterozygous omi^df^/+ (n=15). (D, E) Immunofluorescence images of the apical region of testes stained with (red) anti-Vasa and (green) anti-Bam antibodies from male *c587Gal4, Gal80^ts^* flies that had an RNAi transgene targeting *Nrx-IV* shifted to 30°C for (D) 0 days as control (n=4) or (E) 7 days (n=5) to induce NrxIV knockdown. (F-H) Immunofluorescence images of the apical region of testes stained with (red) anti-Vasa and (green) anti-TJ antibodies from male *c587Gal4, Gal80^ts^* flies shifted to 30°C for 7 days that had an RNAi transgene targeting *side-V* with (F) wild-type *omi* alleles (n=12/15, representative image of 12 shown), (G) heterozygous *omi^df^*/+ (n=6/9, representative image of 6 shown), or (H) heterozygous *omi^df^*/+ (n=3/9, representative image of 3 shown). (I) Quantification of the percentage of *c587Gal4, Gal80^ts^* (Control, n=13), *Nrx-IV* RNAi (n=16), *Nrx-IV* RNAi with heterozygous *omi^df^*/+ (n=16), *side-V* RNAi (n=15), or *side-V* RNAi with heterozygous *omi^df^*/+ (n=9) testes with more TJ-positive cyst cells than Vasa-positive germ cells (Tj > Vasa, light green), equal numbers of TJ-positive cyst cells and Vasa-positive germ cells (Tj = Vasa, teal), and more Vasa-positive germ cells than TJ-positive cyst cells (Vasa > Tj, dark blue). Arrow: Bam expressing germ cells distal to tip of testis. Scale bars: 50 μm.

In contrast to loss of *Nrx-IV*, spermatogonial cell death caused by loss of *side-V* in cyst cells was not substantially rescued in *omi/+* heterozygotes. Compared to knock down of *side-V* in cyst cells in an otherwise wild-type background, knock down of *side-V* in cyst cells in *omi*/+ testes did not result in a significant increase in the number of germ cells (Figure 7F-I).

## Discussion

Our results suggest that the early stages of spermatogonial cyst differentiation involve communication and coordination between germ cells and somatic cyst cells that ensures that once germ cells begin differentiating, septate junctions form between the associated cyst cells to seal the germ cells off into a privileged environment promoting germ cell survival. In wild type, soon after early spermatogonia turn on the differentiation factor *bam*, the two cyst cells that enclose those spermatogonia form septate junctions that set up a permeability barrier (Fairchild et al, 2015) (Figure 8A). Our finding that in *bam* mutant males, where the germ cells maintain an early spermatogonial state (Harris et al., 2025), cyst cells that envelop the overproliferating early germ cells did not seal (Figure 8B) suggests that formation of the septate junctions between cyst cells may be triggered by a signal from the differentiating germ cells they enclose. Stage specific tight junctional complexes form between somatic Sertoli cells in mammalian testes as well, with the resulting Blood Testis Barrier (BTB) leaving proliferating spermatogonia accessible to bodily fluids but isolating the differentiating meiotic and postmeiotic germ cells into a biochemically specialized environment (Mruk & Cheng, 2011). In *c-Kit* mutant mice where the germline fails to differentiate the BTB also does not form (Li et al., 2018), suggesting that similar signals from the germline that trigger occluding junction formation in overlying somatic cells may exist in mammalian spermatogenesis.

**Figure 8.**
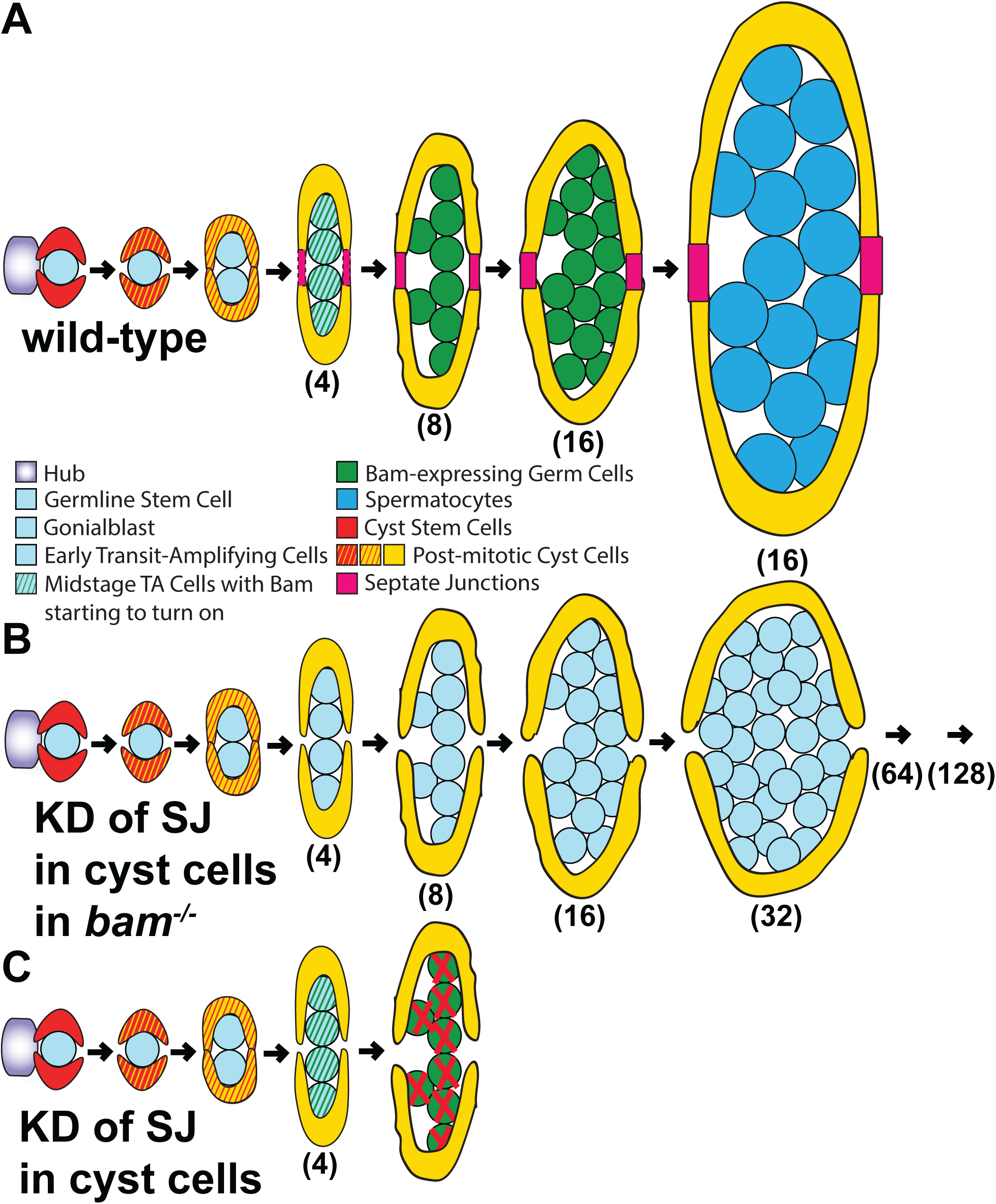
Model of SJ function in cyst cells during early stages of spermatogenesis. (A) Diagram of early stages of *Drosophila* spermatogenesis in wild-type testes highlight the timing of Septate Junction formation coinciding with the onset of Bam expression. (B) Diagram of early stages of *Drosophila* spermatogenesis in *bam*^−/−-^ testes where a Septate Junction component (i.e. *Nrx-IV*) was knocked down in cyst cells, resulting in overproliferation of undifferentiated germ cells. (C) Diagram of early stages of *Drosophila* spermatogenesis in testes where a Septate Junction component (i.e. *Nrx-IV*) was knocked down in cyst cells, depicting the germ cell death observed at the 8-cell stage.

Our data show that once *Drosophila* <u>spermatogonia</u> have begun to express *bam*, if their accompanying cyst cells lack septate junction components, the differentiating germ cells die (Figure 8C). The germ cell death was rescued by loss of function of *bam*, suggesting that while early spermatogonia do not need to be sealed off from the tissue environment, if the two enveloping cyst cells have not established a seal by the time *bam* protein reaches a critical threshold, a surveillance system that detects whether the cyst is sealed off from the tissue environment by functional occluding junctions triggers germ cell death.

The close timing of the initial trigger instructing cyst cells to form septate junctions and the subsequent surveillance mechanism may underlie the longstanding observation that 20-30% of transit amplifying germ cell cysts are eliminated during normal homeostasis (Yacobi-Sharon et al., 2013). The death of some late transit amplifying cysts that occurs in wild type may prune out cases where formation of an effective tight junctional seal by cyst cells did not happen quickly enough to keep up with progression of germ cell differentiation. This pruning of differentiating cysts did not involve the canonical caspase based cell death pathway (Florentin & Arama, 2012) but depended on a mitochondrial protease, *omi* (Yacobi-Sharon et al., 2013), homolog of human HtrA2 implicated in Parkinson’s disease (Plun-Favreau et al., 2007). Consistent with the possibility that failure to form a functional seal between the enclosing cyst cells contributes to the normal elimination of a fraction of late spermatogonial cysts in wild type, lowering function of *omi* suppressed the germ cell death induced by loss of the septate junction component *Nrx-IV* in somatic cyst cells.

The germ cell death resulting from failure to form septate junctions may occur because differentiating late stage spermatogonia cannot tolerate exposure to circulating signaling molecules, peptide hormones or metabolites, although stem cells, gonialblasts and early proliferating spermatogonia such as those that fill *bam* mutant testes can. Alternatively, differentiating late spermatogonia and later meiotic and post meiotic germ cells may require a special microenvironment isolated from bodily fluids and possibly bathed in support factors provided by cyst cells, similar to meiotic and postmeiotic stages in mammalian testes (Xiao et al., 2025).

While the genes required for formation of septate junctions have been investigated extensively in *Drosophila* early embryonic development (Baumgartner et al., 1996; Ward et al., 1998), much remains unknown about the initial molecular events that promote septate junction formation. In *Drosophila* embryos, SJ proteins are expressed before embryonic stage 12, yet they do not form stable complexes until mid-to-late stage 13 (Oshima & Fehon, 2011). The initiating factor for SJ complex formation in embryos has yet to be identified. In *Drosophila* testes, Nrx-IV protein is expressed in both cyst stem cells and differentiating cyst cells, while functional septate junctions able to occlude passage of 10 kDa Dextran into cysts were not established until cysts had reached the 4-8 germ cell stage (Fairchild et al., 2015). Following from our snRNAseq analysis, it is possible that Atpα is a limiting factor, with transcriptional upregulation in cyst cells triggering initiation of septate junction formation. Furthermore, similar bidirectional communication and cross check mechanisms may coordinate formation and sealing of epithelial barriers with differentiation of underlying cells and eliminate assemblies that did not achieve the required arrangement in time for successful differentiation and function in other tissues.

The ability to form septate junctions between cyst cells was previously implicated in differentiation of transit amplifying cells (Fairchild et al., 2015), but not in germ cell survival as shown in this manuscript. However, our result that reducing components of the germ cell death pathway (*omi*^+/−^ flies) rescued the germ cell death caused by knocking down expression of *Nrx-IV* in cyst cells but did not result in germ cell differentiation is consistent with the Fairchild *et al*. (2015) finding. *Nrx-IV* in cyst cells thus likely has at least two roles, required both for survival of germ cells that have embarked on the differentiation program triggered by *bam* and, if the germ cells escape death, for differentiation of spermatogonia into spermatocytes.

Our results underscore the utility of using stage specific gene expression changes revealed in analysis of existing scRNA or snRNA seq data from tissues containing cells in a differentiation sequence to guide small-scale, targeted functional screens. This approach could be particularly useful for identifying functional genes in rare cell types within a tissue. It may also be useful for probing how gene expression changes in one cell type can affect differentiation of partner cells of different origin within tissues, for example changes in epithelial cells affecting the state of underlying mesenchyme in an organ or vice versa. While we did not detect phenotypes in RNAi crosses for many of the genes that were upregulated as cyst cells differentiate, this could be in part due to failure of the RNAi lines tested to knock down mRNAs strongly enough to reveal gene function (Dietzl et al., 2007; Perkins et al., 2015).

One of the genes our screen identified as required in cyst cells for germline survival, *side-V*, is a member of the Side family, a group of proteins that contain Immunoglobulin (Ig) domains and act as receptors for Beaten Path ligands (Zinn & Ozkan, 2017). Loss of function of *side-V* in cyst cells resulted in death of differentiating spermatogonia. This death was rescued in *bam* mutants suggesting that, similar to loss of function of SJ components in cyst cells, it is onset of differentiation that makes germ cells sensitive to lack of *side-V* in somatic cyst cells. However, unlike for loss of *Nrx-IV*, the germ cell death caused by loss of *side-V* function in cysts cells was not substantially rescued by lowering the dose of *omi,* suggesting that the *side-V* dependent germ cell death occurs via a different pathway, or that *side-V* is required at a different step in the sequence of events leading to germ cell death. Both the Beaten Path ligands and the Side receptors are critical for accurate projection of motor neuron axons into muscles (Li et al., 2017; Sink et al., 2001). As members of the Beaten Path family are expressed in both cyst cells and early germ cells in *Drosophila*, they are good candidates for further investigation.

## Acknowledgements

We are grateful to the Bloomington *Drosophila* Stock Center (NIH P40OD018537) and the Vienna *Drosophila* RNAi Center for fly stocks; the Developmental Studies Hybridoma Bank for antibodies; Jaclyn Lim for preliminary experiments involving *Nrx-IV*; and members of the Fuller laboratory for helpful discussions and input on the manuscript. We thank Guy Tanentzapf and Michael Fairchild for productive conversations and feedback on the cell death phenotype following SJ knockdown. CWB was supported by a Stanford Graduate Fellowship and NIH T32AR007422 (PI: Dr. Paul Khavari). This work was supported by NIH grants 1R01GM124054 and R35GM136433 and funds from the Katharine D. McCormick and Stanley McCormick Memorial Professorship and the Reed-Hodgson Professorship in Human Biology to MTF. This work was enabled by the Cell Sciences Imaging Facility (RRID SCR_017787) in the Beckman Center at Stanford University with imaging on the Leica SP8 confocal: Award Number 1S10OD010580-01A1 from the National Center for Research Resources (NCRR).

---

**Supplemental Figure 1.**
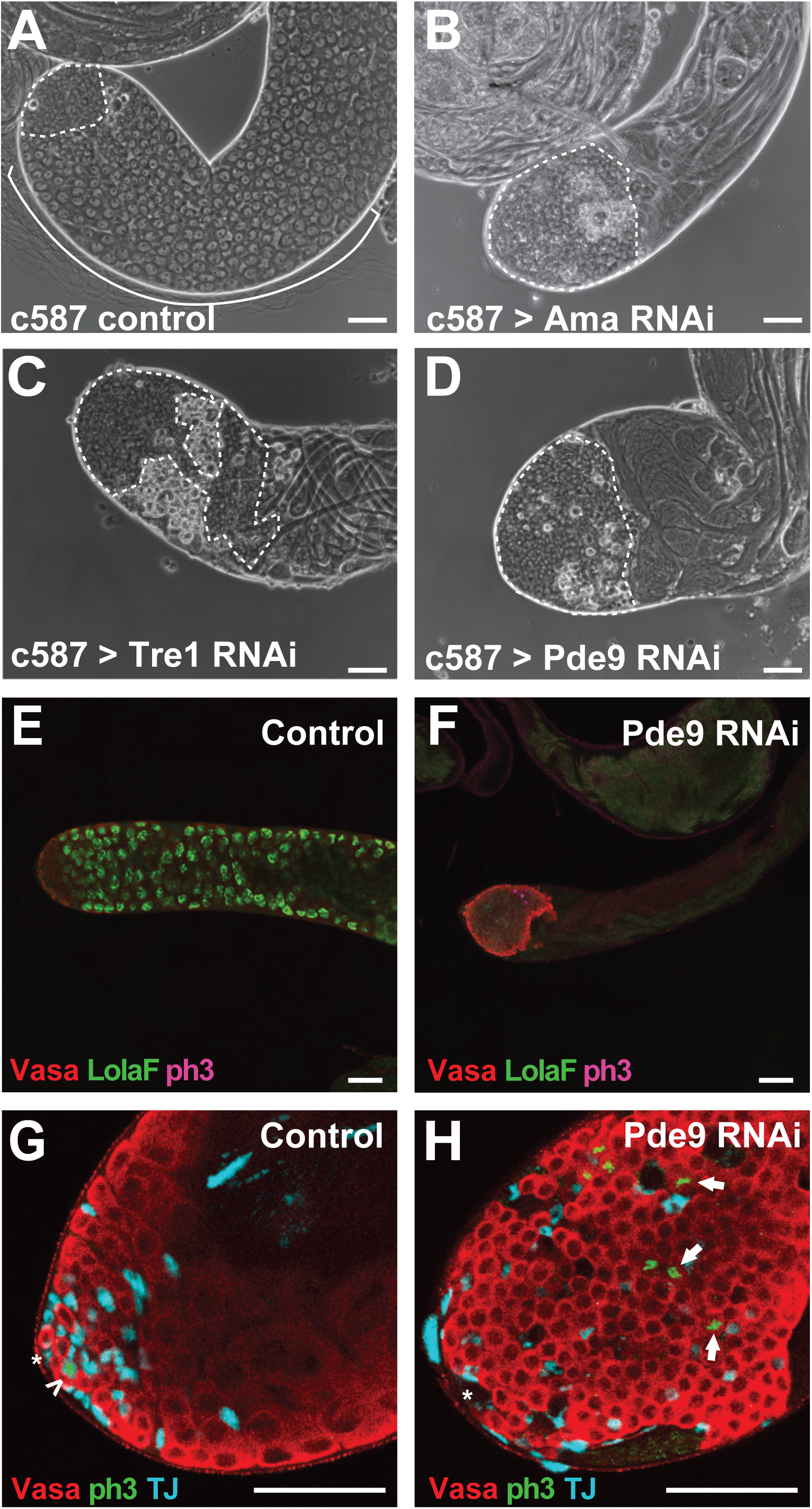
Cyst cell RNAi screen of differentially expressed transcripts reveals novel genes required in cyst cells for differentiation of proliferating spermatogonia. (A-D) Phase-contrast images of the apical region of testes from male *c587Gal4, Gal80^ts^* flies shifted to 30°C for 7 days with (A) no RNAi transgene (n=8) or an RNAi transgene targeting (B) *Ama* (n=8), (C) *Tre1* (n=9), or (D) *Pde9* (n=10). (E, F) Immunofluorescence images of the apical region of testes stained with (red) anti-Vasa and (green) anti-LolaF and (pink) anti-ph3 antibodies from male *c587Gal4, Gal80^ts^*flies shifted to 30°C for 7 days that had (E) no RNAi transgene (n=10) or (F) an RNAi transgene targeting *Pde9* (n=10). (G, H) Immunofluorescence images stained with (red) anti-Vasa, (green) anti-PH3 and (teal) anti-TJ from *c587Gal4, Gal80^ts^* male flies shifted to 30°C for 7 days that had (G) no RNAi transgene (n=9) or a (H) *Pde9* RNAi transgene (n=12). Dotted outline: spermatogonia, curved bracket: spermatocytes, asterisk: hub; arrowhead: dividing spermatogonia near hub; arrows: dividing spermatogonia distal to hub. Scale bars: 50 μm.

**Supplemental Figure 2.**
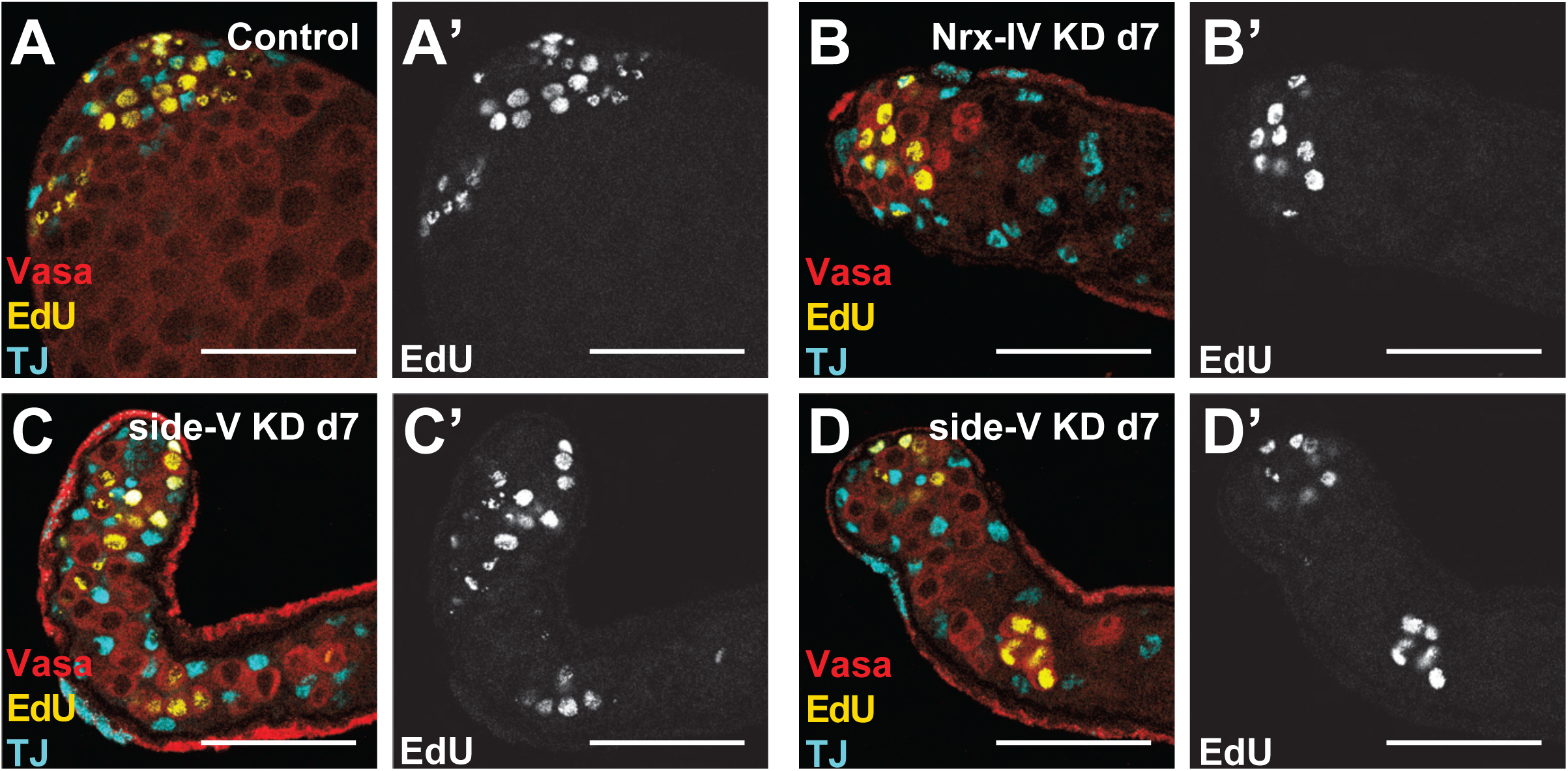
Knockdown of *side-V* in cyst cells triggers loss of germ cells at the 4-cell or early 8-cell stage. (A-D) Immunofluorescence images of the apical region of testes stained with (red) anti-Vasa, (teal) anti-TJ, and (yellow/white) fluorescently-labeled EdU incorporated into cells in S phase from male *c587Gal4, Gal80^ts^* flies shifted to 30°C for 7 days that had (A) no RNAi transgene (n=13), (B) an RNAi transgene targeting *Nrx-IV* (n=11), or (C, D) an RNAi transgene targeting *side-V* (n=10). Scale bars: 50 μm.

**Supplemental Figure 3.**
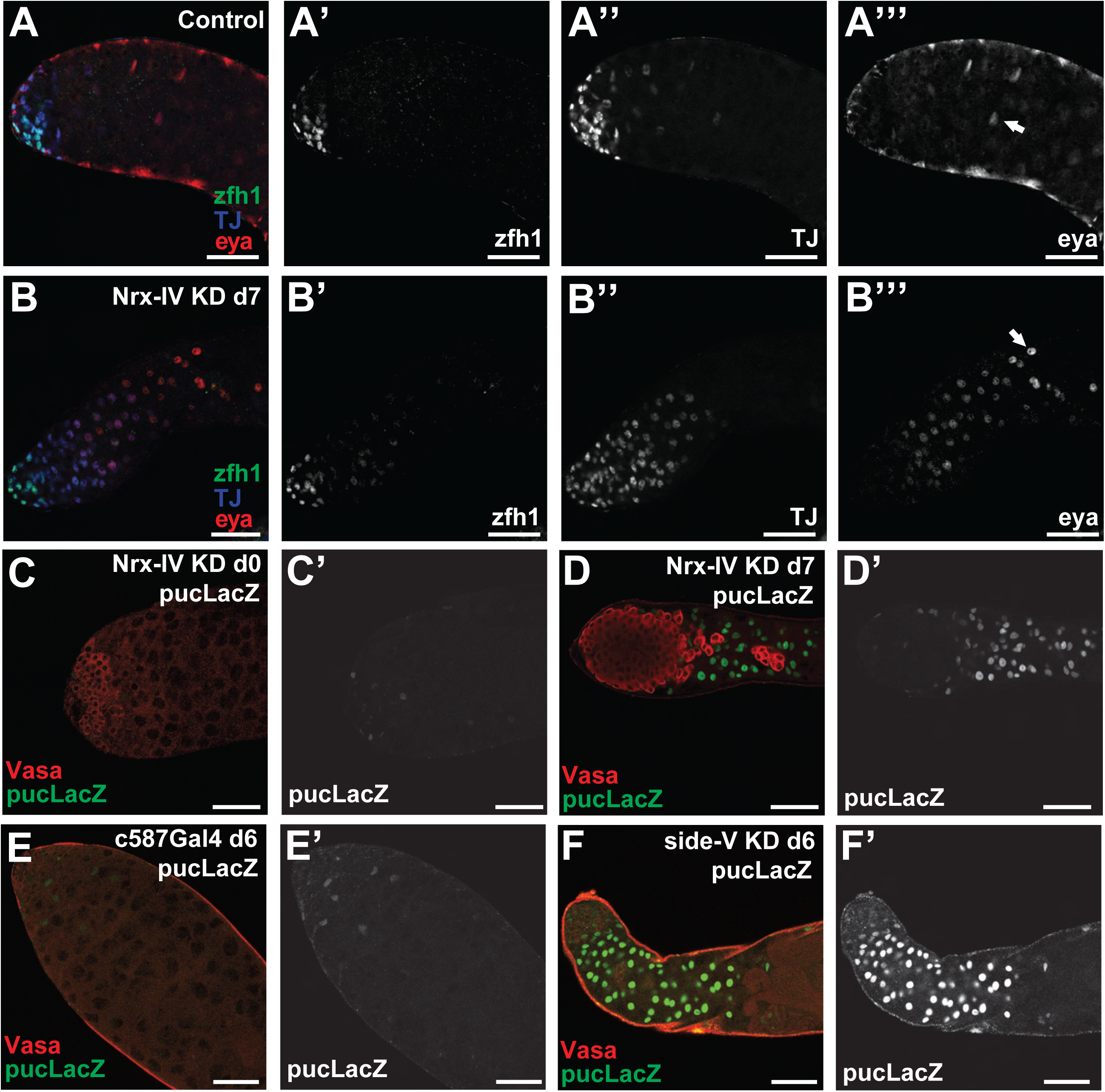
Cyst cells lacking *Nrx-IV* can differentiate, and cyst cells lacking *Nrx-IV* or *side-V* have increased JNK pathway activation. (A, B) Immunofluorescence images of the apical region of testes stained with (green) anti-Zfh1, (blue) anti-TJ, and (red) anti-Eya antibodies from male *c587Gal4, Gal80^ts^* flies shifted to 30°C for 7 days that had (A) no RNAi transgene (n=6) or (B) an RNAi transgene targeting *Nrx-IV* (n=5). (C, D) Immunofluorescence images of the apical region of testes stained with (red) anti-Vasa and (green/white) anti-LacZ antibodies from male *c587Gal4, Gal80^ts^* flies that had a *pucLacZ* transgene and an RNAi transgene targeting *Nrx-IV* shifted to 30°C for (C) 0 days as control (n=8) or (D) 7 days (n=8) to induce knockdown. (E, F) Immunofluorescence images of the apical region of testes stained with (red) anti-Vasa and (green/white) anti-LacZ antibodies from male *c587Gal4, Gal80^ts^* flies that had a *pucLacZ* transgene and were shifted to 30°C for 6 days to induce expression of (E) no RNAi transgene (n=5) or (F) an RNAi transgene targeting *side-V* (n=10). Arrows: differentiated cyst cell. Scale bars: 50 μm.

**Table 1.** Results of targeted RNAi screen in cyst cells using c587Gal4 with temperature sensitive Gal80. Excel spreadsheet indicating testis phenotype for the 255 RNAi lines crossed to *c587Gal4, Gal80^ts^* from male flies shifted to 30°C for 7 days. Includes information on gene name, ID, FBgn, log_2_FC from differential expression analysis of cluster 62 vs cluster 36, VDRC or Bloomington line number, whether a phenotype was observed, and description of phenotype observed.

## Methods

### Fly strains and husbandry

*Drosophila* strains were raised in standard molasses medium at 25°C unless otherwise noted. The following *Drosophila* strains were used: *w^1118^* as wild-type; *c587-Gal80^ts^*; *pucLacZ* (Glise et al., 1995); RNAi fly stocks were obtained from the Vienna *Drosophila* Resource Center (VDRC) (Dietzl et al., 2007) or Bloomington *Drosophila* Stock Center (Bloomington) and further detailed in Table S1. In addition to VDRC lines listed in Table S1, the following lines were used for RNAi knockdown: VDRC8353 (*Neurexin-IV* RNAi), VDRC9788 (*coracle* RNAi), and VDRC107450 (*lachesin* RNAi). In addition to Bloomington lines listed in Table S1, the following line was used for visualization of cyst cell membranes in Figure 6: *UAS-mCD8::GFP* (BDSC5137).

### RNAi knockdown protocol

RNA interference (RNAi) knock-down (KD) experiments were carried out by crossing males with a *UAS-RNAi hairpin* transgene (VDRC or Bloomington) to virgin females of *c587Gal4;ScO/CyO;Gal80^ts^*. Crosses were set up at 22°C and young male progeny were shifted to 30°C to allow inhibition of *Gal80* and expression of the RNAi hairpin for the indicated number of days.

### Immunofluorescence staining

For whole mount staining, testes were dissected from male flies in 1X phosphate-buffered saline (PBS) and incubated with 4% formaldehyde for 20 minutes at room temperature. After fixation, the testes were washed once in PBST (PBS with 0.1% Triton X-100) and permeabilized by incubation with PBS with 0.3% Triton X-100 and 0.6% sodium deoxycholate for 30 minutes at room temperature. After permeabilization, testes were washed once in PBST and blocked for 30 minutes with PBST with 3% bovine serum albumin (BSA), then incubated overnight at 4°C with desired primary antibodies in PBST with 3% BSA. After overnight incubation in primary antibody, testes were washed three times with PBST, incubated with secondary antibodies conjugated with Alexa fluorophores (Alexa Fluor-488, -568, -647 from Molecular Probes) or conjugated with DyLight fluorophores (−594 from Jackson ImmunoResearch Laboratories) in PBST with 3% BSA for 2 hours at room temperature in the dark while rocking, then washed three times in PBST and mounted on glass slides with mounting medium with 1.5 ug/mL DAPI (VECTASHIELD, Vector Labs, Cat# H-1200).

The sources and dilutions of primary antibodies used were as follows: anti-Vasa (goat, 1:100; Santa Cruz Biotechnology, Cat# dc-13); anti-TJ (guinea pig, 1:10,000, a gift from D. Godt, University of Toronto, Canada); anti-Eya (mouse, 1:200, 10H6, DSHB); anti-Bam (mouse, 1:10, DSHB); anti-GFP (chicken, 1:10,000, Abcam 13970); anti-pH3 (rabbit, 1:100, Millipore or mouse, 1:100, ThermoFisher MA5-15220); anti-βGal (mouse, 1:400, Promega Z3781); anti-Hts (mouse, 1:10, 1B1, DSHB); anti-cindr (rabbit, 1:200, a gift from Kaisa Haglund); anti-Fas3 (7G10; 1:10); anti-LolaF (mouse; 1:100; DSHB 1F1-1D5). TUNEL staining used the In Situ Cell Death Detection Kit, TMR Red (Sigma/Roche). Due to the discontinuation of the Santa Cruz Biotechnology anti-Vasa polyclonal antibody during the course of this project, a different anti-Vasa (rat, 1:10, DSHB) primary antibody was used to conduct the stainings in the following figure panels: Figure 2K-L, Figure 3I-P, Figure 5E-F, I-J, M-N, Figure 7F-H, Supplemental Figure 2A-D, Supplemental Figure 3E-F. All remaining panels used the previously noted anti-Vasa primary antibody.

Cells in S phase were labeled with the Click-iT EdU Cell Proliferation Kit for Imaging – Alexa Fluor 555 dye (Invitrogen C10338). Testes were dissected from male flies in Schneider’s (S2) medium and transferred into a drop of the same medium on a slide. The medium was then removed and replaced with 100 μM 5-ethynyl-2’-deoxyuridine (EdU) in S2 cell medium and the testes incubated for 5 minutes at room temperature. Testes were then washed twice in S2 medium and immediately fixed and permeabilized as described above. After permeabilization, testes were washed once in PBST before adding the EdU detection reaction mix per the manufacturer’s instructions. Testes were then incubated in the dark with the reaction mix for 30 minutes, the reaction mix was removed, and the testes were washed twice in PBST. Blocking and antibody protocols continued as described above.

The permeability assay was conducted as described by Fairchild et al. (2015) with the either 10 kDa or 3 kDa AlexaFluor 680 Dextran (ThermoFisher D34681) used. Briefly, testes were dissected from males in Schneider’s (S2) medium and transferred into a drop of 10 kDa or 3 kDa Dextran in S2 medium at a concentration of 0.2 μg/μL. After addition of the cover slip, testes were imaged live using a confocal microscope (see below).

### Fluorescence microscopy and image analysis

Fluorescent images were taken with a Leica SP8 confocal. Laser intensity and detector gain for the confocal microscope or exposure time were adjusted for each experiment to ensure that the signal was in linear range and not saturated. Once image acquisition settings were determined, the same settings were maintained for the entire set of images be compared. Immunofluorescence images were processed using ImageJ with all samples to be compared going through the same processing.

### ASAP analysis of differentially expressed transcripts

To identify transcripts upregulated at the earliest stages of cyst cell differentiation the ASAP analysis program was accessed at https://asap.epfl.ch/ and used to run differential expression analysis between cluster 62 (cyst stem cells) and 36 (early cyst cells) of Leiden resolution 6.0 of the testis snRNAseq dataset (Li et al., 2022). The parameters used were the Wilcoxin [Seurat] method of differential expression, minimum % cells > 0.1, and fold change threshold of 1.3.

